# From museum drawer to tree: historical DNA phylogenomics clarifies the systematics of rare dung beetles (Coleoptera: Scarabaeinae) from museum collections

**DOI:** 10.1101/2023.10.27.564347

**Authors:** Fernando Lopes, Nicole Gunter, Conrad P.D.T. Gillett, Giulio Montanaro, Michele Rossini, Federica Losacco, Gimo M. Daniel, Nicolas Straube, Sergei Tarasov

## Abstract

Although several methods exist for extracting and sequencing historical DNA originating from dry-preserved insect specimens deposited in natural history museums, no consensus exists as to what is the optimal approach. We demonstrate that a customized, low-cost archival DNA extraction protocol (∼ €10 per sample), in combination with Ultraconserved Elements (UCEs), is an effective tool for insect phylogenomic studies. We successfully tested our approach by sequencing DNA from scarab dung beetles preserved in both wet and dry collections, including unique primary type and rare historical specimens from internationally important natural history museums in London, Paris and Helsinki. The focal specimens comprise enigmatic dung beetle genera (*Nesosisyphus, Onychotechus* and *Helictopleurus*) that varied in age and preservation. The oldest specimen, the holotype of the now possibly extinct Mauritian endemic *Nesosisyphus rotundatus*, was collected in 1944. We obtained high-quality DNA from all studied specimens to enable the generation of a UCE-based dataset that revealed an insightful and well-supported phylogenetic tree of dung beetles. The resulting phylogeny suggested the reclassification of *Onychotechus* (previously *incertae sedis*) within the tribe Coprini. Our approach demonstrates the feasibility and effectiveness of combining DNA data from historic and recent museum specimens to provide novel insights. The proposed archival DNA protocol is available at DOI 10.17504/protocols.io.81wgbybqyvpk/v1

**Highlights:**

- We combined custom low-cost archival DNA extractions and Ultraconserved Element phylogenomics
- DNA from rare museum specimens of enigmatic dung beetles revealed their phylogenetic connections
- Genomic data was obtained from the holotype of a potentially extinct monoinsular endemic species
- Genomic data allowed a rare and enigmatic species of previously unknown affinity to be classified
- The morphology of museum specimens remained intact following non-destructive DNA extraction

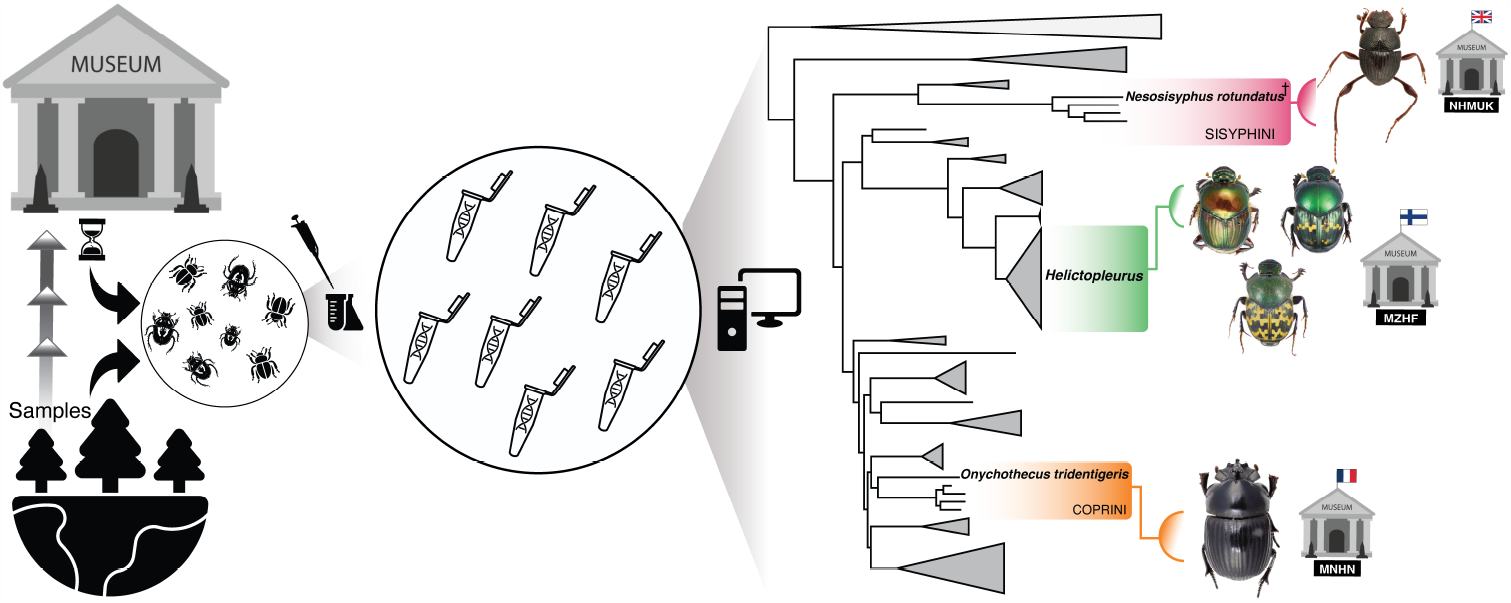

## 1. Introduction

Museomics, a term encompassing procedures allowing access to and analysis of the historical genomic data preserved in biological specimens deposited in natural history museums, is providing unprecedented opportunities to investigate evolutionary histories (Miller et al., 2009; Raxworthy and Smith, 2021). Together with concomitant advances in high-throughput sequencing technologies and bioinformatics, museomics has paved the way for the exploitation of an ever-broader diversity of taxonomic and temporal sampling (Orlando et al., 2015; Burrell et al., 2015). Importantly, by enabling access to genomes already preserved in existing museum specimens, museomics can circumvent the need for costly, laborious, and unpredictable bespoke fieldwork, in order to achieve taxon sampling objectives (Raxworthy and Smith, 2021; Orlando et al., 2015). Museomics is also compatible with physically preserving the morphological integrity of specimens, when non-destructive DNA extraction methods are employed. This is of paramount importance to natural history museums and the scientific community because it ensures that intact voucher specimens will remain available for study by future generations (Fong et al., 2023; Weirauch et al., 2020). Indeed, the importance of museomics can only heighten as the necessity for inclusion of recently extinct species within phylogenies becomes increasingly inevitable (Toussaint et al., 2021). Progress in insect museomics has already greatly contributed to the study of insects - Earth’s most diverse organisms (Eggleton, 2020; Fong et al., 2023). However, despite notable achievements, challenges still remain (Raxworthy and Smith, 2021). Specifically, DNA in many dry-preserved museum specimens is fragmented and prone to contamination, whilst the comparatively small amount of tissue present in small insects can further limit the success of DNA extractions (Patzold et al., 2020).

In recent years, a variety of molecular methods have been developed to obtain historical DNA data at a genome-wide scale (Orlando et al., 2015; Burrell et al., 2015), including approaches exploiting both whole-genome (shotgun) and reduced-genome sequencing (Jin et al., 2020; Toussaint et al., 2021; Twort et al., 2021). Many widely used methods rely on standard DNA extractions using commercial DNA kits, followed by the construction of DNA libraries based on hybridization approaches that combine restriction enzyme fragmentation and RNA probe capture. For instance, HyRAD uses a double enzymatic restriction of DNA extracts from fresh samples (containing well-preserved DNA) to produce RNA probes that serve as baits for capturing homologous fragments from historical (more degraded) DNA libraries (Suchan et al., 2022; Gauthier et al., 2020). However, standard DNA extraction, typically undertaken with commercially available kits, is optimized for high molecular weight DNA, only ineffectively capturing low-weight short fragments, which are precisely those expected from degraded historical samples. Furthermore, reduced genome approaches exploiting restriction enzymes require a comparatively large initial amount of source DNA, not easily obtained from small insect specimens (Sproul and Maddison, 2017). Moreover, those methods tend to be costly when extensively sampling a wide range of insect taxa. They are also labor-intensive because they require the creation of custom RNA probes for each taxon being studied. Crucially, such methods are susceptible to the drawbacks associated with restriction enzymes. These include the potential for enzyme mismatch, either due to point mutations because target taxa are too distantly related or due to DNA fragmentation at restriction sites (especially in poorly preserved samples), both possibilities that can lead to missing data (Cerca et al., 2021).

In phylogenetic studies with historical DNA, cost-effective methods such as Ultra Conserved Elements (UCEs) and Anchored Hybrid Enrichment (AHE) are rising in popularity due to their ability to target specific informative loci within a focal group (Faircloth, 2017; Call et al., 2021; Faircloth et al., 2012; Mayer et al., 2021). While standard extraction from dry-preserved specimens may yield adequate DNA for UCE and AHE sequencing (Gustafson et al., 2020; Mayer et al., 2021), its success varies based on preservation. Therefore, exploring more sensitive extraction methods is essential, especially since their application to entomological collections remains poorly investigated (Call et al., 2021).

In this study, we aim to bridge this gap by assessing a cost-effective (∼ €10 per sample) archival DNA extraction protocol (Straube et al., 2021) tailored for historical insect specimens, specifically for UCE-sequencing. We applied this protocol to explore the phylogenetic relationships of eleven species and subspecies of dung beetles (Coleoptera: Scarabaeinae) represented by historical specimens from three museums: The Natural History Museum, London (NHMUK); the Muséum National d’Histoire Naturelle, Paris (MNHN); and the Finnish Museum of Natural History, Helsinki (MZHF).

The specimens are of diverse ages and represent enigmatic species of questionable phylogenetic assignment (Table 1). The oldest specimen, the holotype from NHMUK of *Nesosisyphus rotundatus* collected in 1944, is a potentially extinct species from Mauritius, not previously included in molecular phylogenies. The extremely rare Oriental genus *Onychothecus*, with uncertain taxonomic affinity (Tarasov and Dimitrov, 2016) and previously lacking DNA data, was represented by a specimen from MNHN, collected in 1985. Finally, nine poorly-known taxa belonging to the endemic Madagascan genus *Helictopleurus* were represented by specimens from MZHF collected between 2003–2010. Our archival DNA extraction protocol yielded high quality DNA for successful UCE-sequencing using the recently designed probe set for scarab beetles (Gustafson et al., 2023). To elucidate the phylogenetic position of the selected enigmatic species, we expanded our taxon sampling by sequencing additional alcohol-preserved dung beetles using a standard DNA extraction protocol with a commercial kit. In the following sections, we discuss the phylogenetic placement of the focal species based on our results and provide necessary taxonomic changes. We also explore the broader application of the proposed extraction protocol to a wide range of historical specimens of insects and other taxa.

**Table 1.** Dry-preserved scarab dung beetle specimens from natural history museum collections, used in historical DNA extractions. Natural History Museum, London (NHMUK); Muséum National d’Histoire Naturelle, Paris (MNHN); and Finnish Museum of Natural History (MZHF).

| Species | Sampling Year | Museum | Origin |
| --- | --- | --- | --- |
| *Nesosysiphus rotundatus* Vinson, 1946 [holotype] | 1944 | NHMUK | Mauritius |
| *Onychothecus tridentigeris* Zelenka, 1992 | 1985 | MNHN | Thailand |
| *Helictopleurus fasciolatus fasciolatus* (Fairmaire, 1898) | 2003 | MZHF | Madagascar |
| *Helictopleurus sinuatocornis* (Fairmaire, 1898) | 2003 | MZHF | Madagascar |
| *Helictopleurus perrieri* (Fairmaire, 1898) | 2006 | MZHF | Madagascar |
| *Helictopleurus undatus* (Olivier, 1789) | 2008 | MZHF | Madagascar |
| *Helictopleurus nicolleti* Lebis, 1960 | 2008 | MZHF | Madagascar |
| *Helictopleurus near furcicornis* Lebis, 1960 | 2008 | MZHF | Madagascar |
| *Helictopleurus neuter* (Fairmaire, 1898) | 2009 | MZHF | Madagascar |
| *Helictopleurus fasciolatus obscurus* Lebis, 1960 | 2009 | MZHF | Madagascar |
| *Helictopleurus fasciolatus pseudofasciolatus* Montreuil, 2007 | 2010 | MZHF | Madagascar |

## 2. Material and methods

### 2.1 Taxon sampling

We compiled a dataset of UCE sequences from 96 beetles (Table S1), encompassing mostly scarabaeoid beetle lineages from various biogeographical regions. Our dataset combined 70 newly-sequenced specimens for this study with existing data for 26 specimens from a previous study and available on GenBank (Gustafson et al., 2023). The ingroup consisted of 67 samples belonging to 42 genera or subgenera of true dung beetles of the subfamily Scarabaeinae. The outgroup consisted of 29 samples (of 26 genera) belonging either to scarab beetle families and subfamilies other than Scarabaeinae, or to non-scarab beetles (two species of Silphidae). 59 of the newly sequenced samples (representing 40 genera) originated from frozen (-20°C), alcohol-preserved “wet collection” specimens that were sourced from five natural history museums. A further 11 historical samples belonging to three genera (*Nesosisyphus, Onychotechus* and *Helictopleurus*) were selected from the dry collections of three museums, and formed the focal taxa in our study (Table 1, Fig. 1A).

**Figure 1:**
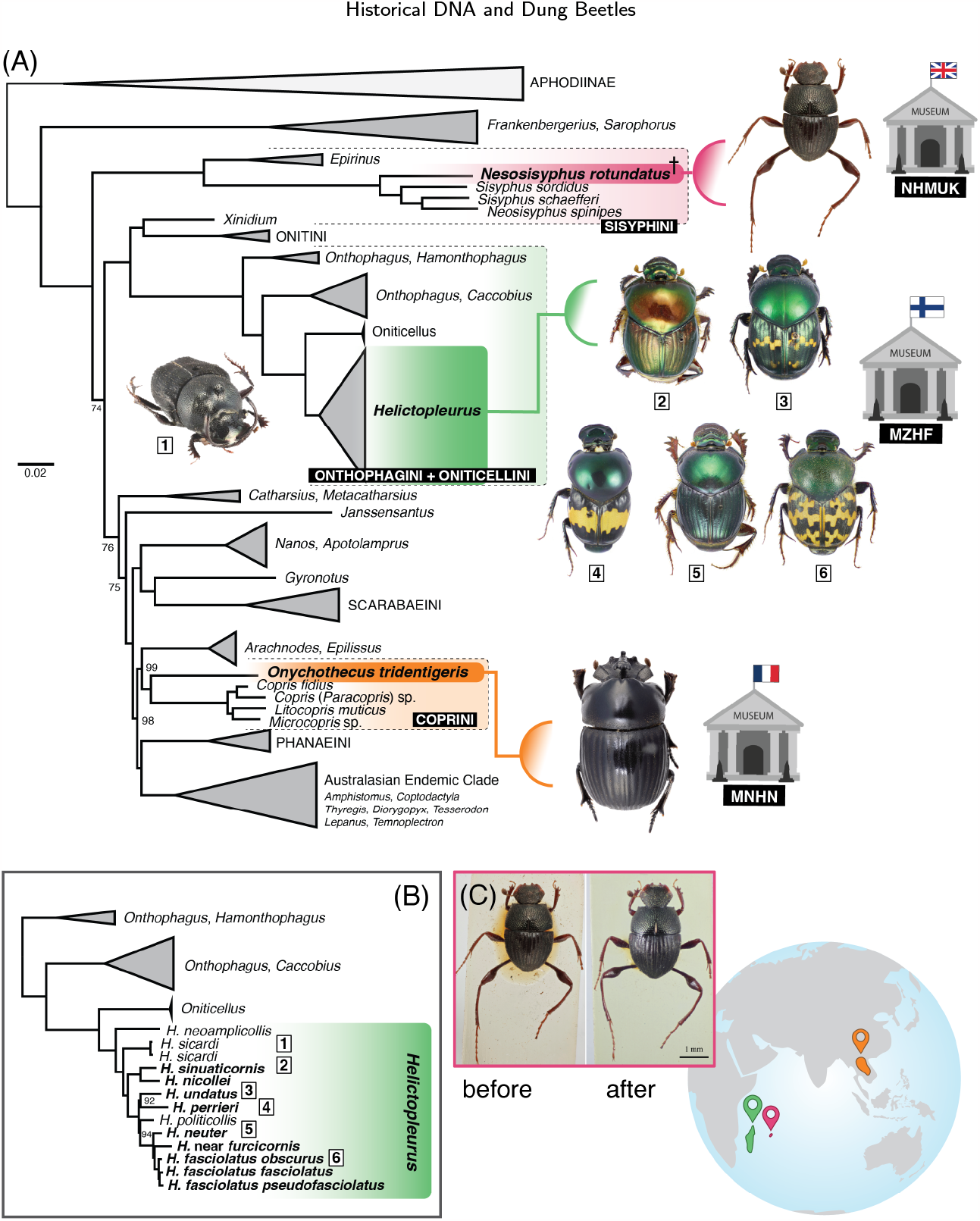
(A) Species tree inferred using 1,497 UCEs and the 50% complete dataset; dry-preserved historical museum specimens are in bold. (B) The part of the tree showing the relationships within *Helictopleurus*. (C) The holotype of *Neosisyphus rotundatus* before and after DNA extraction using the proposed archival protocol. Node numbers in (A) and (B) indicate bootstrap values < 100, while all other nodes have full bootstrap support.

### 2.2 DNA extraction: archival DNA protocol

Our optimized protocol is available at Protocols.io. Briefly, the protocol is a customization of the archival DNA extraction protocol and Guanidine treatment described by Straube et al. (2021) which, in turn, was influenced by the studies of Dabney et al. (2013) and Rohland et al. (2004). The new approach was first proposed for wet-preserved vertebrates and is based on the binding of DNA to a PCR purification silica membrane in the presence of a chaotropic salt (guanidine hydrochloride) buffer (Table S2 and at Protocols.io). The method uses an extension reservoir attached to a commercial silica spin column, able to retain DNA fragments of lengths varying from 70 bp to 4 kbp. This adaptation allows for a more than tenfold increase in the ratio of binding buffer to sample and enhances the recovery of short DNA fragments, typically present in historical samples (Straube et al., 2021). We further customized this protocol as a non-destructive extraction protocol using dry-preserved beetle specimens from several museum entomological collections, as specified in step 4 of the protocol. In short, we optimized the way we used the samples for the lysis step by not destroying body parts. DNA extractions from dry-preserved museum specimens (Table S1) were undertaken in a dedicated ‘clean room’ for historical samples at MZHF. *Nesosysiphus rotundatus*, a monoinsular endemic species om Mauritius that is known only from two specimens, was epresented by its holotype, deposited in NMHUK (Fig. 1C). The tiny specimen, one of the smallest scarab dung beetles in the world (∼4 mm), was collected by J. Vinson in 1944 (Vinson, 1946). This specimen was carefully relaxed and disarticulated; only the prothorax (with exposed internal tissues) with attached forelegs (but not the head) was used as a source of DNA during the digestion step of the extraction (Fig. 1C), resulting in a total of 17.03 ng of DNA that generated 1,528 UCE loci after sequencing. Following DNA extraction, the digested body parts remained well-preserved, without visible external deterioration, and the specimen was successfully reassembled (Fig. 1C). *Onychothecus tridentigeris* is a much larger, very rare species from Thailand, that was represented by a non-type specimen (∼20 mm) deposited in MNHN (Table 1, Fig. 2). From this specimen, we destructively used an entire leg during the extraction (Fig. 2B,C), which resulted in 17.40 ng of DNA that generated 1,692 UCE loci.

**Figure 2:**
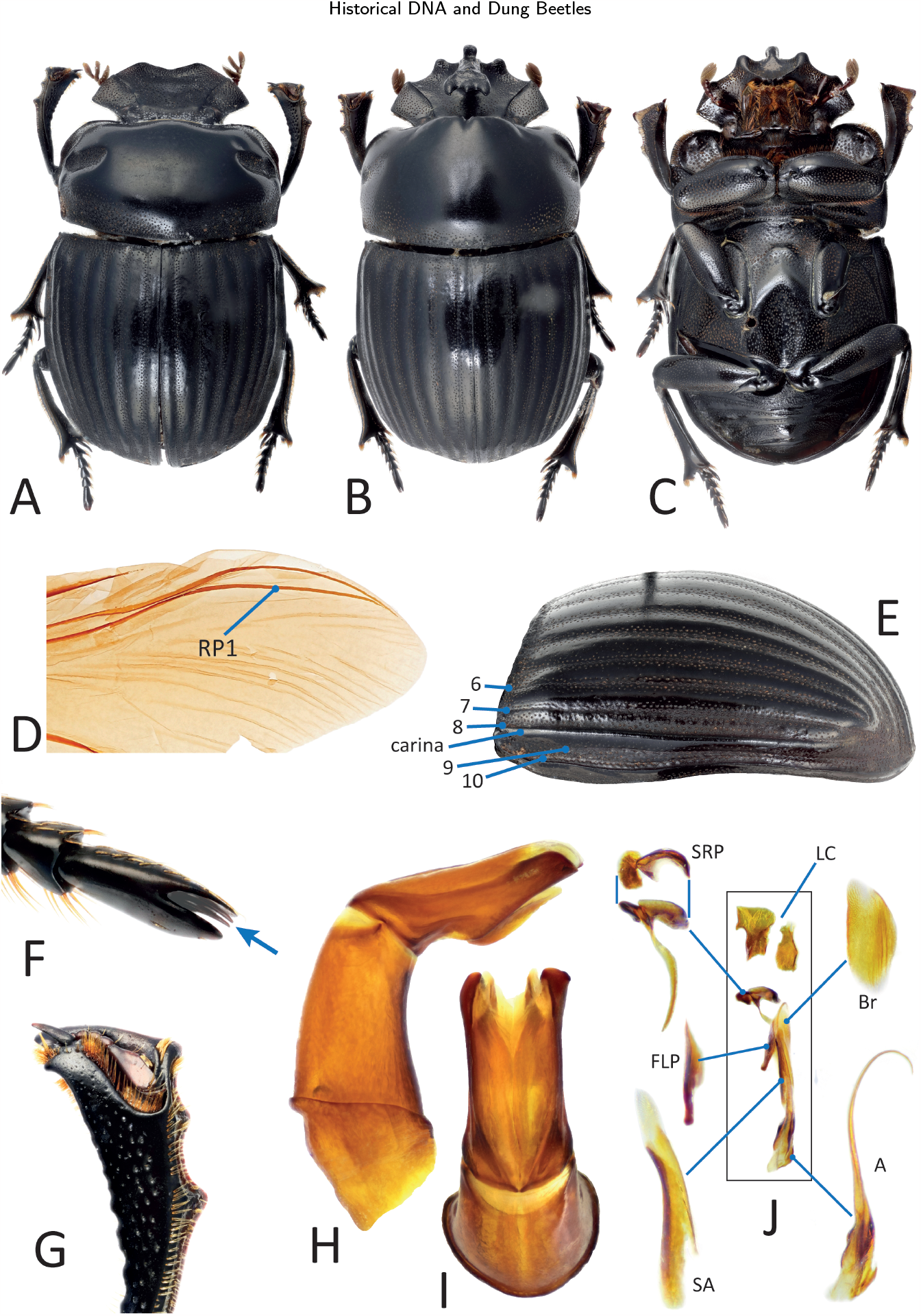
Morphology of *Onychothecus tridentigeris* Zelenka, 1992. Dorsal habitus of male (A) and female (B); ventral view of female (C); right wing in dorsal view, with radial posterior vein 1 (RP1) indicated (D); left elytron in lateral view, with elytral striae numbered and lateral carina (E); hind tarsus, with the modified terminal tarsomere concealing the claws (F); right protibia of male, in dorsal view (G); aedeagus in lateral (H) and dorsal (I) views; endophallites (J) (abbreviations follow Tarasov and Génier (2015)).

### 2.3 DNA extraction: alcohol-preserved specimens

DNA was extracted from wet-preserved museum specimens using the QIAamp DNA Micro Kit (QIAGEN) following the manufacturer’s protocol. 2 *μ*l of the resulting purified DNA from both dry-preserved (historical) and wet-preserved specimens was quantified on a Qubit fluorometer 4.0 (Thermo Fisher Scientific) using high-sensitivity reagents.

### 2.4 Library preparation and sequencing

We used the tailored UCE Scarabaeinae probe-set Scarab 3Kv1 (Gustafson et al., 2023), a genome reduction approach for beetle genomes combining the Coleoptera 1.1Kv1 probeset (Faircloth, 2017) and Scarabaeinae-specific targeted loci (Gustafson et al., 2023), to generate sequence data for reconstruction of phylogenies (Fig. 1 and Figs. S1-S4). This probe set contains 25,786 probes targeting 3,174 loci (Gustafson et al., 2023). Library preparation and enrichment were performed at RAPiD Genomics LLC (Gainesville, FL, U.S.A.) for Illumina sequencing, applying their high-throughput workflow with proprietary chemistry. DNA was sheared to a mean fragment length of 500 bp and A-tailed, followed by the incorporation of unique dual-indexed Illumina adaptors and PCR enrichment. Libraries were sequenced on an Illumina HiSeq 2500 (2 × 150 pairend).

### 2.5 Data Processing

Demultiplexing and trimming were performed by RAPiD Genomics LLC using Illumina bcl2fastq2 2.20 (Illumina, 2017). Our UCE datasets were assembled using the package Phyluce 1.7.3 (Faircloth, 2016) following our workflow available on GitHub. Raw demultiplexed reads were first cleaned using Illumiprocessor 2.0 (Faircloth, 2013) to remove residual adapter contamination. Cleaned reads were inspected for quality using FastQC (Andrews et al., 2010) and assembled into contigs in Spades 3.15.4 (Bankevich et al., 2012). The Scarab 3kv1 UCE probes were matched to the assembled contigs in Phyluce, with a minimum identity of 80% and coverage of 80× to avoid off-target contaminating sequences (Gustafson et al., 2023; Bossert and Danforth, 2018). UCE loci were then extracted from the sequenced data. We harvested UCE loci from the available whole beetle genomes on GenBank (Gustafson et al., 2023) (Table S1) using Phyluce and combined them with our newly sequenced data. The UCE loci were aligned in MAFFT (Katoh and Standley, 2013), using the default Phyluce settings, and the command *-no-trim* to provide internal trimming, as recommended for analysis of divergences over 50 million years old (Faircloth, 2016). The resulting alignments were parsed to a parallel wrapper around Gblocks to eliminate poorly aligned positions and divergent regions using the settings: b1=0.5, b2=0.85, b3=8, b4=10 (Castresana, 2000; Talavera and Castresana, 2007). Summary statistics for the generated datasets were computed using the program AMAS (Borowiec, 2016).

### 2.6 Phylogenomics

For phylogenomic analyses, we constructed data matrices for species trees and Maximum-Likelihood (ML) concatenated-based inferences with 50% and 70% complete data, allowing up to 50% and 30% missing taxa for each locus, respectively (Molloy and Warnow, 2018; Gustafson et al., 2020). Hereafter, the 50% and 70% complete datasets will be called 50p and 70p datasets. Full and partitioned UCE alignments are available on the Open Science Framework (OSF) repository at DOI 10.17605/osf.io/mxwj7. ID labels in tree files were translated into full species names using the custom Python script *rename_leaves_v1*.*0*.*py* available on GitHub.

#### 2.6.1 Species tree analysis

We recovered phylogenies accommodating for gene tree heterogeneity due to Incomplete Lineage Sorting (Maddison, 1997; Edwards, 2009) considering each UCE loci as an independent gene. Species trees were estimated in IQ-Tree 2.0.7 (Minh et al., 2020), with confidence levels calculated using 1000 ultrafast bootstrap (UFBoot) replicates, and topologies tested by the Shimodaira–Hasegawa test (SH-aLRT). The best substitution models were automatically selected using ModelFinder (*-m mfp* option) implemented in IQ-Tree under the Bayesian Information Criterion (Kalyaanamoorthy et al., 2017).

To reduce the risk of overestimating branch support with UFBoot owing to severe model violations, we used a hill-climbing nearest-neighbor interchange (NNI, *-bnni* option) topology search strategy to optimize each bootstrap tree. As phylogenetic models rely on various simplifying assumptions to ease computations (*e.g*., treelikeness, reversibility, and homogeneity of substitution models), estimations of some genomic regions can severely violate model assumptions, causing biases in phylogenetic estimates of tree topologies (Naser-Khdour et al., 2019). To test these violations on each locus, we also applied the test of symmetry with the option *-*-*symtest-remove-bad*. Partitions (concatenated analyses) and genes (species trees analyses) with *p*value *≤* 0.05 for the test of symmetry were removed from downstream analyses (Naser-Khdour et al., 2019).

#### 2.6.2 Concatenated-based analysis

We also constructed ML phylogenies from the concatenated 50p and 70p datasets using IQ-Tree, using the same parameters mentioned above for species trees.

The datasets were partitioned with Sliding-Window Site Characteristics (SWSC-EN), an entropy-based method developed specifically for UCE data (Tagliacollo and Lanfear, 2018). To implement the SWSC-EN method, we used Phyluce to generate a concatenated *Nexus* file with the location of each UCE locus as character sets. With the SWSC-EN Python 3.6 script, configuration files were created to be used with Partitionfinder 2.1.1 and Python 2.7 (Lanfear et al., 2017). As Partitionfinder2 works only with *Phylip* alignments, we converted the concatenated *Nexus* file to Relaxed *Phylip* format using Geneious 2022.2.1 (Kearse et al., 2012). The partitioning scheme was then generated with Partitionfinder2 with linked branch lengths, a GTR+G model of evolution, an Akaike information criterion with correction (AICc) for model selection, and a variant of the relaxed hierarchical clustering search algorithm (supplementary data) (Lanfear et al., 2014; Gustafson et al., 2020).

### 2.7 Morphological Examination

As a complement to molecular inference of the phylogenetic position of *Onychothecus* and related taxa, we studied the morphology of two available specimens of *Onychothecus tridentigeris* (deposited in MNHN) in detail. Morphological terminology and protocols follow Tarasov and Dimitrov (2016) and Tarasov and Génier (2015). Specimens were examined under a Leica S9D stereomicroscope. Photographs were taken with a Canon MP-E 65 mm, f/2.8, 1–5× macro lens mounted on a Canon EOS 5D camera, and then stacked using the Stackshot (Cognisys Inc.) automated system.

## 3. Results

### 3.1 UCE data

We obtained a mean of 1.86×10^7^paired-end reads per sample. Our results revealed that shorter fragments from museum samples were effectively integrated into DNA libraries, resulting in the recovery of a substantial number of UCE loci for downstream phylogenomic analyses (Fig. 3, Table S1). Specifically, samples for which DNA was extracted using the historical DNA protocol yielded the highest number of recovered loci (2,264), followed by alcohol-preserved samples extracted using the commercial kit (1,620 loci) and UCE data retrieved from GenBank genomes (909 loci; Table S1 and Fig. 3B). Interestingly, older samples, such as *O. tridentigeris* and *N. rotundatus*, exhibited a similar number of recovered loci compared to the more recently collected wet-preserved samples extracted with the commercial kit. (Table 1 and Table S1).

Before data filtering, the full concatenated alignment (96 tips) contained 3,160 UCE loci and 269,808 parsimony-informative sites distributed across 675,2 Kbp (Table S3). The 50p dataset contained 1,497 UCE loci with a mean of 79.97 parsimony-informative sites per locus (Table S4), and the 70p dataset contained 289 UCE loci with a mean of 60.54 parsimony-informative sites per locus (Table S5). The conspicuous disparity among unfiltered, 50p and 70p datasets is due to large proportion of missing data present in the genomes retrieved from GenBank, which mostly served as outgroup taxa in our study (Gustafson et al. (2023) Table S2 and Figs. S1-S5).

**Figure 3:**
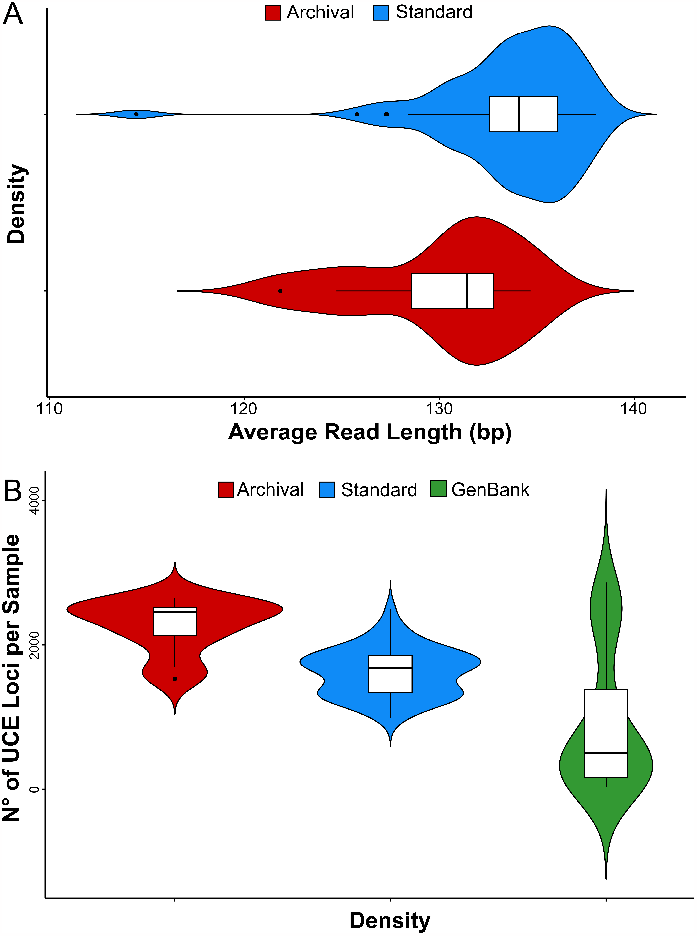
Summary of UCE data resulting from two DNA extraction methods (archival extraction protocol in red and standard extraction in blue) and beetle genomes from GenBank (in green, only in B). Violin plots illustrate the kernel density and boxplots display the median and variation. (A) Distribution of read length generated per sample, demonstrating that the density of shorter reads was generally higher from archival extractions. (B) Distribution of the number of UCE loci per sample, demonstrating that the greatest density of samples that generated large numbers of captured loci resulted from archival extractions.

### 3.2 Phylogenomics

Increasing the completeness threshold during the construction of data matrices significantly decreased the number of UCE loci and overall bootstrap support (Fig. 1, Figs. S1-S4 and Table S4–S5). Because phylogenomic studies generally do not benefit from filtering out loci with an increased proportion of missing data (Molloy and Warnow, 2018), we focused on the 50p dataset, which resulted in an optimal trade-off between the highest overall bootstrap support and SH-values (Fig. 1 and Figs. S1-S2), and the number of recovered loci (Table S1).

Our phylogenetic trees based on the concatenated alignment (ML tree) were generally well-supported, with only a few nodes of moderate depth having poor support (see Fig. 1 and Figs. S1-S2). Notably, Scarabaeinae formed a monophyletic group, with *Frankenbergerius* and *Sarophorus* representing a basal lineages which is sister to the remaining Scarabaeinae. The Afrotropical *Epirinus* was identified as the sister taxon to the remaining Sisyphini. Additionally, the Australasian genera appeared to form a monophyletic group, as too did the clade consisting of the tribes Onthophagini and Oniticellini.

The oldest historical specimen of *N. rotundatus* from Mauritius consistently clustered within the tribe Sisyphini across all analyses with strong bootstrap support (BS: 100; Fig. 1A and C). The enigmatic species, *O. tridentigeris* from the Oriental Region, consistently emerged as the sister taxon to the genera of the tribe Coprini in all trees (BS: ≥ 98; Fig. 1A and Fig. 2). All Madagascan *Helictopleurus* species formed a sister clade to Oniticellini in all analyses as well (Figs. 1A and B). Some slight variations in the grouping of certain *Helictopleurus* species were observed in the 70p datasets, likely due to reduced genomic information and/or short branches in the backbone of the clade (Fig. 1 and S1-S4).

## 4. Discussion

### 4.1 Phylogenetic relationships

All resultant topologies were generally consistent with previous morphological (Tarasov and Génier, 2015) and molecular analyses based on individual genes (Tarasov and Génier, 2015; Tarasov and Dimitrov, 2016; Gunter et al., 2016) and UCE data (Gustafson et al. (2023), but containing only limited taxon sampling).

The present phylogeny, whilst encompassing a relatively taxonomically diverse set of dung beetles, is not comprehensive enough to justify discussion of general phylogenetic relationships and their taxonomic consequences within Scarabaeinae. Therefore, we leave that broader discussion to forthcoming phylogenomic studies that will incorporate substantially larger taxon sampling. Instead, we focus on assessing the relationships among the enigmatic taxa herein sequenced from historical specimens.

#### Mauritian *Nesosisyphus*

This study offers the first insights into the relationships of the endemic Mauritian genus *Nesosisyphus* that has not previously undergone phylogenetic analysis. Our placement of *N. rotundatus* (Fig. 1A, C) within the tribe Sisyphini (Daniel et al., 2020; Tarasov and Dimitrov, 2016) was expected and is also supported by morphological synapomorphies that clearly indicate that *Nesosisyphus* is a member of this tribe. Therefore, the phylogenetic result based on UCE data obtained from the old holotype of *N. rotundatus* strongly agrees with our initial hypothesis (Vinson, 1946).

*Nesosisyphus rotundatus* is a flightless roller dung beetle, uniquely known by the two specimens that make up the type series collected during the early 1940s from the southern slope of Mount Ory in Mauritius (Vinson, 1946). All subsequent collecting efforts on the island, which have included sampling at the type locality, and have resulted in the discovery of three additional Mauritian endemic species of *Nesosisyphus* (Losacco et al., *in prep*.), failed to relocate this species. Given this, and considering the rapid loss of indigenous habitats and biodiversity in Mauritius, in general, due to anthropogenic habitat destruction and the introduction of exotic species (Safford, 1997; Monty et al., 2013), we regard this species as potentially extinct (Losacco et al., *in prep*). One of the achievements of our study has been to unlock genomic data from this enigmatic species for further investigation.

#### Oriental *Onychothecus*

This extremely rare genus comprises four species distributed in southeastern Asia (Figs. 1 and 2): China (Yunnan), Myanmar, Thailand, Laos, and Vietnam (Ochi and Kon, 1998; Schoolmeesters, 2023). It is remarkable for displaying secondary sexual dimorphism that is unusual within scarab beetles; females bear a cephalic horn and males are hornless (the reverse is overwhelmingly more common in the superfamily Scarabaeoidea), in addition to having an unknown diet, habits, and general biology. *Onychothecus* has not yet been classified (=*incertae sedis*) into any of the existing tribes in the subfamily Scarabaeinae (Tarasov and Dimitrov, 2016). Only a single previous phylogenetic treatment of the genus exists (Montreuil, 1998), based upon morphological data, which identified it as a sister to the genus *Paraphytus* that has a disjunct Afrotropical and Oriental distribution. *Paraphytus* belongs to the most basal lineage of Scarabainae (Tarasov and Dimitrov, 2016) that also includes the Afrotropical genera *Frankenbergerius* and *Sarophorus*, which we included in the present analyses. Our resulting phylogeny indicates that *Onychothecus* does not belong to that basal lineage, being instead recovered as sister to the clade containing the genera *Copris, Litocopris, Microcopris*, belonging to the tribe Coprini *sensu* Tarasov and Dimitrov (2016). Consequently, based on our results, we assign *Onychothecus* to the tribe Coprini and discuss this placement in a separate section below.

#### Madagascan *Helictopleurus*

The genus *Helictopleurus* comprises approximately 65 species, all endemic to Madagascar (Figs. 1A-B) and primarily occurring in forest habitats (Wirta et al., 2010; Miraldo et al., 2011). We extracted and sequenced DNA from nine dry-preserved specimens from museum collections, having been collected between 2003 and 2010, in addition to two alcohol-preserved specimens. Sequence data for one additional species was included from a previously published study (Rossini et al., 2021). Our phylogenetic analyses (Figs. 1A-B, S1-S4) confirm previous results, demonstrating the monophyly of the genus *Helictopleurus*, its sister relationship to the genus *Oniticellus*, and that the *Helictopleurus* + *Oniticellus* clade falls within the clade containing Onthophagini + Oniticellini (Breeschoten et al., 2016; Tarasov and Solodovnikov, 2011; Wirta et al., 2008). However, when compared to earlier molecular phylogenies based on individual genes, our analyses resulted in slight variations in the interspecific relationships within *Helictopleurus* (Rossini et al., 2021; Wirta et al., 2008). Enhancing taxon sampling in future studies and potentially integrating it with prior single-gene data will help achieve more robust results.

The fact that our results, including newly sequenced data from museum specimens of varying ages (Table 1) and preservation, produced robust results that are consistent with existing phylogenies (Gustafson et al., 2023; Tarasov and Dimitrov, 2016), demonstrates the effectiveness of the proposed historical DNA approach in combination with UCE sequencing. Such consistency is of particular significance because concerns about sequence data obtained from historical specimens being contaminated or of poor quality, and consequently obfuscating or impeding phylogenetic inference, appear not to have been borne out in our study.

### 4.2 Tribal transfer of *Onychothecus* to Coprini

#### Genus *Onychothecus* Boucomont, 1912

- *Onychothecus* Boucomont, 1912: original description; *as member of* Scatonomini Lacordaire, 1856 *synonym of* Deltochilini Lacordaire, 1856: *sensu* Bouchard et al. (2011).
- *Onychothecus*; Balthasar (1963): *as member of* Pinotini Kolbe, 1905 *synonym of* Ateuchini Perty, 1830: *sensu* Smith (2006).
- *Onychothecus*; Tarasov and Dimitrov (2016): *as incerta sedis*.
- **Type species:** *Onychothecus ateuchoides* Boucomont, 1912.

The tribe Coprini is distributed in both the Old and New World and includes five genera: *Copris, Litocopris, Microcopris, Pseudocopris*, and *Pseudopedaria*. The tribe lacks unique apomorphies allowing for unequivocal diagnosis. Instead it is characterized only by a combination of six characters (Tarasov and Dimitrov, 2016).

We examined the morphology of two specimens of *Onychothecus tridentigeris* in detail, including the first-known male of the species (Figs. 2A and 2G-J). Our phylogenetic analyses strongly support the position of *Onychothecus* as a sister taxon to a clade containing the five genera making up the Coprini. Based upon this evidence, two alternative taxonomic actions were be considered: either creating a new tribe to accommodate *Onychothecus* or assigning it to Coprini. We have chosen the latter option and, herein, treat the genus *Onychothecus* as a member of the tribe Coprini. In our opinion, this action is justified in order to maintain stability in the classification of Scarabaeinae.

Although the general habitus of *Onychothecus* resembles that of many other members of the tribe Coprini, its morphology stands out within the Scarabaeinae in general. Specifically, *Onychothecus* exhibits dorsally excavated protibial apices, terminal tarsomeres that conceal the tarsal claws and, most strikingly, ‘inverse sexual dimorphism’, wherein the female is the horned sex. Additionally, it possesses characters that have not previously been adopted to diagnose Coprini, including laterally carinate elytra, an absence of the posterior sclerite of the wing, and an absence of the posterior ridge of the hypomera. These observations therefore oblige revision and expansion of the morphological diagnosis of the tribe Coprini. We present the new diagnosis of Coprini in Table 2.

**Table 2.** Updated diagnosis of the tribe Coprini. The combination of characters 1–6 constitutes a diagnosis of Coprini that includes *Onychothecus*; for details see Tarasov and Dimitrov (2016). Characters 7–10 (marked with ^*^) refer to autapomorphies of *Onychothecus*.

| Character | *Onychothecus* | other Coprini genera |
| --- | --- | --- |
| 1. Wing apex, posterior sclerite | absent (Fig. 2D) | present |
| 2. Number of elytral striae | 10 (Fig. 2E) | 10 |
| 3. Elytral stria 8 | carinate (Fig. 2E) | not carinate |
| 4. Superior right peripheral (SRP) endophallite | not ring-shaped (Fig. 2J) | not ring-shaped |
| 5. Hypomera, anterior ridge reaches lateral margin | present | present |
| 6. Posterior longitudinal hypomeral ridge | absent | usually present |
| *7. Last tarsomere | concealing tarsal claws (Fig. 2F-G) | not concealing tarsal claws |
| *8. Protibial apex | excavated dorsally (Fig. 2G) | not excavated dorsally |
| *9. Parameres | asymmetric (Fig. 2H-I) | usually symmetric |
| *10. Sexual dimorphism | only females with cephalic horn | often males with cephalic horn |

## 4.3 Conclusion

We successfully obtained genomic data that allowed for the phylogenetic placement of several species represented by unique historical specimens deposited in important natural history museum collections (Table 1). Our results demonstrate that combining a minimally destructive and low-cost archival DNA extraction (∼ €10 per sample), with subsequent target enrichment of DNA libraries for sequencing a curated set of beetle UCE loci, is an efficient museomics tool (see Tables S1-S2). The proposed extraction protocol should also integrate well with AHE sequencing. Even from the limited quantity of source tissue available in old museum specimens, by being able to capture small fragments of degraded DNA (Fig. 2), we demonstrated a remarkably favorable trade-off between preserving specimen morphology and generating informative genomic-level data. Our customized extraction protocol can be performed using standard equipment commonly available in molecular laboratories within two days, including an overnight digestion step. The protocol is optimized for 4–6 samples in each extraction batch (see protocol). Because the procedure is designed to capture small amounts of fragmented DNA, prone to cross-contamination, simultaneous handling of a larger number of samples is not recommended, in order to minimize this risk. We believe that the method’s strength is that it is particularly applicable when extractions from old specimens deposited in museum collections are necessary. Because many taxa are rare and known only by unique or very few valuable specimens held in museums, nondestructive museomics methods, such as the one we have described, are essential in order to allow for such (often inordinately interesting) taxa to be included in phylogenomic studies.

## Acknowledgements

We are grateful to the Coleoptera curators of collaborating natural history museums for the loan of rare specimens under their care: Max Barclay (The Natural History Museum, London) and Olivier Montreuil (Muséum national d’Histoire Naturelle, Paris). We thank the Malagasy Institut pour la Conservation des Ecosystèmes Tropicaux (MICET) for their help in acquiring research permits and for their logistical support during fieldwork in Madagascar. We also acknowledge the generous technical support given to us by Louise Lindblom, head of the DNA lab, University Museum of Bergen, in relation to the historical DNA extraction protocol. Finally, from the Finnish Museum of Natural History, Helsinki, we thank all members of the Tarasov lab and the Coleoptera team for their constructive suggestions and discussions, in addition to the staff of the DNA lab, especially the head of the DNA Lab, Gunilla St åhls, for their support.

## Authorship contribution statement

**Fernando Lopes:** Conceptualization of this study, data curation, methodology, investigation, analyses, visualization, writing - original draft preparation. **Nicole Gunter:** Funding acquisition, data curation. **Conrad P.D.T. Gillett:** Resources, data curation, writing - review & editing. **Giulio Montanaro:** Data curation, investigation, writing - original draft preparation. **Michele Rossini:** Data curation. **Federica Losacco:** Data curation, visualization. **Gimo M. Daniel:** Data curation. **Nicolas Straube:** Data curation, writing - review & editing. **Sergei Tarasov:** Funding acquisition, resources, conceptualization of this study, data curation, methodology, investigation, visualization, writing - original draft preparation.

## Data Availability

The data presented in this study can be accessed at Open Science Framework and Protocols.io via the two following hyperlinks: DOI 10.17605/osf.io/mxwj7 and DOI 10.17504/protocols.io.81wgbybqyvpk/v1. Raw sequencing data can be found at NCBI BioProject PRJNA1031114.

